# The extracellular matrix fibulin 7 maintains epidermal stem cell heterogeneity during skin aging

**DOI:** 10.1101/2022.02.28.482241

**Authors:** Erna Raja, Gopakumar Changarathil, Lalhaba Oinam, Yen Xuan Ngo, Jun Tsunezumi, Ryutaro Ishii, Takako Sasaki, Kyoko Imanaka-Yoshida, Hiromi Yanagisawa, Aiko Sada

## Abstract

Tissue stem cells divide infrequently as a protective mechanism against internal and external stresses associated with aging. Here, we demonstrate that slow- and fast-cycling interfollicular epidermal stem cells in mouse skin undergo distinct aging processes. Two years of lineage tracing reveals that Dlx1+ slow-cycling clones expanded into the fast-cycling stem cell territory, while the number of Slc1a3+ fast-cycling clones gradually declined. Transcriptome analysis further indicated that the molecular properties of each stem cell population are altered with age. Mice lacking fibulin 7, an extracellular matrix (ECM), show early impairments resembling epidermal stem cell aging, such as the loss of fast-cycling clones, delayed wound healing, and increased expression of inflammation- and differentiation-related genes. Fibulin 7 interacts with structural ECM and matricellular proteins, and the overexpression of fibulin 7 in primary keratinocytes results in slower proliferation in the absence or presence of inflammatory cytokine IL-6. Thus, these results suggest that fibulin 7 plays a crucial role in maintaining tissue resilience and epidermal stem cell heterogeneity during skin aging.

## Introduction

Stem cells in the skin respond to external and internal factors, such as physical injury, inflammation, and mitotic stress, and activate various cellular functions to promote tissue remodeling. Skin becomes thinner and more fragile as we age, and its ability to recover from damage or stress declines. The skin epithelium can be divided into the interfollicular epidermis (IFE) and its appendages (i.e., hair follicles, sebaceous glands), each of which is maintained by its own stem cell population (Gonzales & Fuchs, 2017). In the IFE, epidermal stem cells in the basal layer divide and differentiate upward during homeostasis to provide barrier functions and repair damage. Hair follicle stem cells (HFSCs), which support hair growth, behave independently of epidermal stem cells under homeostatic conditions, but have the plasticity to switch their fate to the epidermal lineage during skin injury or tumorigenesis (Ge *et al*, 2017; Gonzales *et al*, 2021; Ito *et al*, 2005). In chronologically aged skin, ectopic differentiation, migration, and loss of HFSCs, melanocytes, or epidermal stem cells have been observed (Inomata *et al*, 2009; Liu *et al*, 2019; Matsumura *et al*, 2016; Zhang *et al*, 2021). However, whether skin aging occurs primarily due to stem cell-intrinsic factors or because of changes in the microenvironment or dermal compartment is still debated (Ge *et al*, 2020).

Stem cell adhesion and interaction with the extracellular matrix (ECM) are crucial in self-renewal and fate regulation, and alterations in quality and quantity of ECM can induce aging-associated skin dysfunction (Egbert *et al*, 2014; Ge *et al*., 2020; Koester *et al*, 2021; Liu *et al*., 2019; Watanabe *et al*, 2017; Watt & Fujiwara, 2011). Skin aging leads to DNA damage and proteolysis of collagen XVII, a hemidesmosome component important for HFSC and epidermal stem cell maintenance, proliferation, and polarity (Liu *et al*., 2019; Matsumura *et al*., 2016; Watanabe *et al*, 2021; Watanabe *et al*., 2017). Aging skin is also associated with reduced wound healing ability (Keyes *et al*, 2016; Liu *et al*., 2019) and concomitant with decreased expression of some dermal ECM proteins, including periostin and tenascin C (Choi *et al*, 2020; Egbert *et al*., 2014). However, little is known about the factors that regulate ECM changes in epidermal stem cells during aging.

Tissue stem cells are generally thought to be protected against aging by their less frequent division. Stem cell aging may be caused by repeated replication stress and accumulation of DNA damage (Behrens *et al*, 2014), which culminates in molecular damage, metabolic and epigenetic alterations, aberrant proliferation and differentiation, and depletion of stem cell pools (Ermolaeva *et al*, 2018). The slow-cycling populations of hair follicle (Keyes *et al*, 2013; Lay *et al*, 2016; Wang *et al*, 2016) and hematopoietic (Sacma *et al*, 2019) stem cells have protective mechanisms against aging. Although the slow-cycling nature of stem cells can minimize DNA damage, mutations acquired in slow-cycling stem cells are more likely to accumulate over time due to the error-prone DNA repair pathway non-homologous end joining, in contrast to the homologous recombination that predominantly occurs in faster-cycling stem cells (Tumpel & Rudolph, 2019). Whether this slow-cycling speed serves as a mechanism to maintain long-term stem cell potential and delay stem cell aging in the skin remains unclear.

The slow- and fast-cycling stem cells in the IFE of mouse skin enable us to study the relationship between cell division frequency and aging by directly comparing aging processes within the same tissue environment. Using the H2B-GFP system, we previously reported that slow-cycling (label-retaining cells; LRC) and fast-cycling (non-label-retaining cells; nLRC) populations were molecularly distinct and highly compartmentalized in the IFE of mouse skin (Sada *et al*, 2016) (Fig. 1A, S1A). Slow-cycling stem cells express *Dlx1*, while fast-cycling stem cells express *Slc1a3*. Slow- and fast-cycling stem cells undergo distinct differentiation programs to give rise to the keratin (K)10+ interscale and K31+ scale structures of the tail skin, respectively (Gomez *et al*, 2013; Sada *et al*., 2016). They stay within their territorial boundaries in homeostasis but retain plasticity to contribute to each other’s territory during injury repair (Sada *et al*., 2016). However, it remains unknown how these two stem cell populations age and whether different molecular mechanisms regulate their respective aging processes. Here, we found that aging led to disruption of the stem cell compartment in the IFE, along with a reduction in the number of fast-cycling stem cell clones and a diminished skin resilience to external tissue damage and inflammation. We further identified fibulin 7 as a regulator of the stem cell aging process that controls long-term stem cell potential and ECM maintenance.

**Figure 1.**
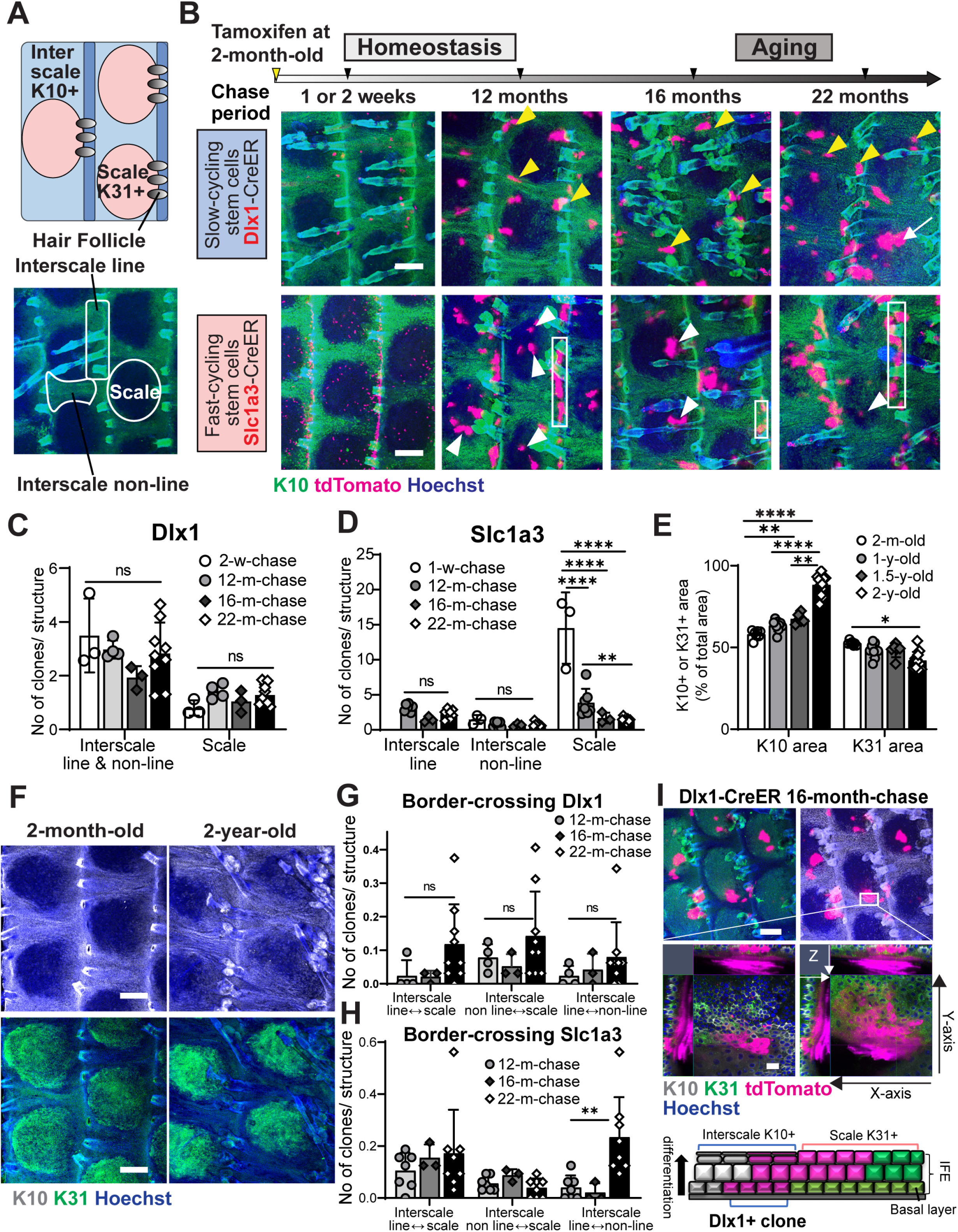
Long-term lineage tracing depicting depletion of fast-cycling stem cell clones and dysregulation of the stem cell compartmentalization in chronological aging. **A.** Upper diagram describes the interscale and scale localizations in the interfollicular epidermis (IFE), marked by keratin (K) 10 and K31, respectively. Lower wholemount tail epidermis image shows delineations of areas defined as scale, interscale line, and interscale non-line. **B.** Low-dose tamoxifen was administered once at 2 months of age to allow single-cell labeling and tail skin samples were collected after 1–2-week-, 12-month-, 16-month-, and 22-month-chases. Wholemount staining of tail epidermis with tdTomato, K10, and Hoechst. Scale bar, 200 μm. Yellow and white arrowheads are examples of Dlx1^CreER+^ interscale and Slc1a3^CreER+^ scale clones, respectively. White boxes indicate examples of Slc1a3^CreER+^ interscale line clones. A white arrow indicates a clone that is crossing boundary. **C.** The number of Dlx1^CreER+^ clones per interscale or scale structure. ns.: not significant. The number of mice and chase time: *N*=3 (2 weeks), *N*=4 (12 months), *N*=3 (16 months), and *N*=9 (22 months). **D.** The number of Slc1a3^CreER+^ clones per interscale or scale structure. ****; *P*<0.0001. **; *P*<0.01. The number of mice and chase time: *N*=3 (1 week), *N*=7 (12 months), *N*=3 (16 months) and *N*=8 (22 months). **E.** Quantitation of K10+ or K31+ areas normalized to total IFE areas. The number of mice at age 2 months *N*=7, 1 year *N*=9, 1.5 years *N*=6, 2 years *N*=12. *; *P*<0.05. **F.** Wholemount staining of K10 and K31 in 2-month-old compared to 2-year-old mice. Scale bar, 200 μm. **G, H.** The number of Dlx1^CreER+^ and Slc1a3^CreER+^ clones crossing the interscale–scale boundaries. The number of animals used is the same as in (C–D). All statistical tests in Figure 1 were performed with two-way ANOVA, Tukey’s multiple comparisons test. All graphs show mean ± S.D. **I.** Confocal imaging of representative clones at 1.5-year-chase, stained with K10 and K31. White box indicates area that is enlarged in the lower panel. Z-stack images show the clone is originating from the basal layer and expanding into the upper differentiated layers. Cartoon summarizes the sagittal view of the clones. Scale bars, 200 μm (upper panel). Scale bars, 20 μm (lower panel). K10 and K31 intensities in (B, F, I) were adjusted to similar levels between samples.

## Results

### Fast-cycling stem cells are gradually lost and compartments of distinct stem cell populations impaired during aging

To address long-term stem cell potential and its behavioral changes during aging, lineage tracing was performed using Dlx1^CreER^ and Slc1a3^CreER^ to mark slow- and fast-cycling stem cells, respectively. As previously observed (Sada *et al*., 2016), from 1-week to 1-year post-labeling during homeostasis, Dlx1 clones were enriched in the interscale (yellow arrowheads, Fig. 1B), whereas Slc1a3 clones were preferentially localized in the scale and interscale line regions (white arrowheads and white boxes, Fig. 1B). We found that the number and localization of Dlx1 clones were less affected during aging (Fig. 1B, C), but the number of Slc1a3 clones in the scale was significantly decreased, starting at a 1-year chase (Fig. 1B, D). It is noteworthy that while Slc1a3-labeled fast-cycling stem cell clones in the scale were gradually lost during aging, the small number of Slc1a3 clones in the interscale continued to thrive and expand within the interscale line and non-line regions (Fig. 1B, D). Hence, fast-cycling stem cell clones were gradually depleted during aging, but slow-cycling stem cell lineages were maintained.

Along with changes at the stem cell level, wholemount images of K10 and K31 staining (the regional differentiation markers of interscale and scale, respectively) demonstrated a significant age-dependent increase in the K10-positive interscale area in 2-year-old mice compared with 2-month-old mice (Fig. 1B, 1E–F). With aging, there was a significant decrease in the number of fast-cycling stem cell clones in the scale, but effects on the size of the K31-positive scale region were minimal (Fig. 1E, F), suggesting a possible change of the slow-cycling stem cell fate to the scale lineage occurring at the interscale-scale boundary. Indeed, a small number of clones were found to cross the borders between the interscale and scale regions over time, and the frequency of these border-crossing clones tended to increase in 2-year-old mice (arrow in Fig. 1B, G, H, S1 B–C). Using higher magnification Z-stack images from Dlx1 aging clones at 16- and 22-month-chases, it became apparent that the cells crossing the interscale-scale borders primarily consisted of upper differentiated cells (expressing both K10 and K31 markers), whereas basal cells were located in the interscale (Fig. 1I, S1D, S1F). In other words, the conversion from slow- to fast-cycling lineage may not be occurring at the stem cell level. At a 22-month-chase, we also found Slc1a3 clones that expanded beyond the boundary from the interscale line region to scale, which may have compensated for the decrease of scale clones (Fig. S1E, S1F). This result suggests that aging alters the unique lineage fate of epidermal stem cells and disrupts their compartmentalization within the tissue.

### Aging induces alterations in molecular properties of slow- and fast-cycling stem cells

To investigate age-dependent molecular changes, bulk RNA-sequencing (RNA-seq) was performed using slow-cycling and fast-cycling IFE populations isolated from the tail skin of 2-month-old and 2-year-old mice (Sada *et al*., 2016; Tumbar *et al*, 2004). Since most fast-cycling clones labeled with Slc1a3^CreER^ in the scale region were depleted with age, making cell isolation difficult, we utilized the K5-tTA/pTRE-H2B-GFP mouse model to isolate a fast-cycling cell population (nLRC) at 2 years of age. Wholemount immunostaining showed differences in GFP dilution rates after doxycycline administration in interscale LRCs and scale nLRCs in both 2- to 3-month-old and 2-year-old mice (Fig. S2A, B). Epidermal basal cells (α6-integrin^high^/CD34^-^/Sca1^+^) from 2-month-old mice that underwent 0–2 cell divisions were defined as LRCs, whereas nLRCs were those that experienced ≥ 4 divisions (Fig. S2A, C, left panel in D). In 2-year-old mice, due to slower basal cell proliferation, LRCs and nLRCs were defined as cells with 0–1 divisions and ≥ 3 divisions, respectively (Fig. S2B, C, right panel in D). The slowest dividing 10–15% and fastest dividing 20–30% of the total basal IFE population were isolated in both age groups (Fig. S2E).

In principal component analysis (PCA), LRCs and nLRCs showed clustering separation, indicating their unique transcriptome identities and age-dependent changes (Fig. 2A, Table S1). We first analyzed aging-induced changes in signature gene expression for young, 2-month-old LRCs (total 407 genes) and nLRCs (total 331 genes) (Fig. 2B). Results showed that the following percentages of these signature genes were altered with age: 23% of young LRC genes and 39% of young nLRC genes were downregulated, whereas 7% of young LRC genes and 6% of young nLRC genes were upregulated in the 2-year-old counterparts (Fig. 2B, Fig. S2F-I and Table S2). Nonetheless, some of the 2-month-old signature gene expression patterns were maintained during aging (Fig. S2J-K, Table S2), suggesting that some characteristics of young LRCs or nLRCs are retained.

**Figure 2.**
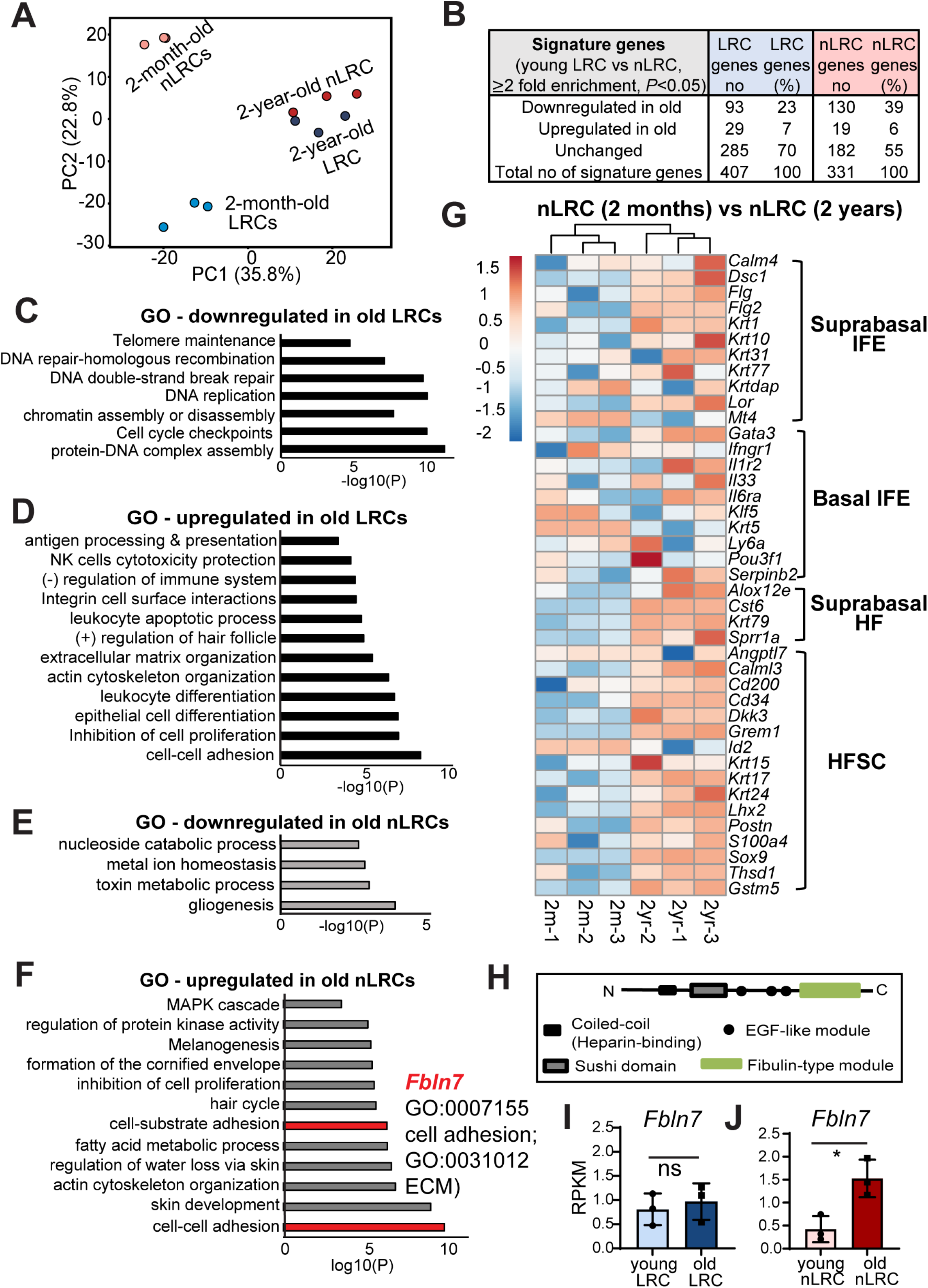
Global gene expression analysis reveals molecular changes in slow- and fast-cycling epidermal stem cells during aging. **A.** PCA map describes the transcriptomic clustering of LRCs and nLRCs constructed from 3-fold differentially expressed genes among 2-month-old (*N*=3) and 2-year-old (*N*=3) mice. **B.** Table summarizes the number of signature genes in young LRCs vs. nLRCs and their respective changes in 2-year-old skin. **C-F.** Gene ontology analysis obtained from ≥2-fold differentially regulated genes (*P*<0.05) in 2-year-old LRCs or nLRCs compared to 2-month-old mice. **G.** Heatmap describes changes in basal and suprabasal signature genes of epidermal stem cells and HFSCs (Ge *et al*., 2020) in 2-month-old vs. 2-year-old nLRCs. Heatmap scale shows log2 RPKM values. **H.** Schematic of fibulin 7 protein structure. **I, J.** *Fbln7* gene expression is increased in aged nLRCs (J) but not in aged LRCs (I). Statistical test: t-test. *N*=3 mice.

To determine molecular factors that contribute to the loss of epidermal stem cell heterogeneity during aging, and particularly the depletion of fast-cycling stem cells, we further interrogated gene expression changes in 2-year-old LRCs and nLRCs. Gene ontology (GO) analysis showed that a group of genes involved in DNA damage repair, telomere maintenance, DNA replication, and chromatin regulation was markedly reduced in old LRCs, suggesting that aged slow-cycling epidermal stem cells may be more prone to accumulating DNA damage (Fig. 2C, Table S1). Moreover, the 2-year-old LRCs upregulated genes related to immune response (Fig. 2D, Table S1). The nLRCs, on the other hand, were characterized by changes in genes related to cell metabolism (Fig. 2E, F, Table S1), which has been implicated in controlling stem cell proliferative heterogeneity and aging (Nakamura-Ishizu *et al*, 2020). GO analysis thus suggests that gene regulation differs depending on the stem cell population. Over-representation of processes that were common to the 2-year-old LRCs or nLRCs versus the young LRCs or nLRCs was also observed, including cell adhesion, ECM and actin cytoskeleton dynamics, inhibition of proliferation, induction of differentiation, and hair follicle development (Fig. 2D, F, Table S1).

To evaluate whether an imbalance between stem cell self-renewal and differentiation contributes to age-related clonal decline in fast-cycling epidermal stem cells, we examined changes in the expression patterns of established epidermal lineage marker genes (Ge *et al*., 2020). This analysis showed that the expression of IFE differentiation genes and hair follicle-lineage markers, which are repressed in young nLRCs, were enhanced in older ones, suggesting that the maintenance of undifferentiated status and lineage identity may be compromised in aged fast-cycling epidermal stem cells (Fig. 2G). Thus, the RNA-seq results and lineage tracing analysis suggest that epidermal stem cells with different division frequencies undergo distinct cellular and molecular processes and show functional decline with aging.

### Loss of *Fbln7* accelerates age-dependent depletion of fast-cycling stem cell clones and delays wound healing

Because both microenvironment and stem cell-intrinsic mechanisms govern the stem cell aging process, we focused on the secreted ECM genes of 2-year-old fast-cycling stem cells, whose stem cell potential/lineage is compromised in aged skin. Among the genes upregulated in the 2-year-old nLRCs was fibulin 7 (gene symbol *Fbln7*), a secreted glycoprotein belonging to the short fibulin family of ECM proteins (Fig. 2F, H-J). Fibulin 7 regulates cell differentiation and migration through interaction with other ECM proteins, heparin, and cell surface receptors such as β1 integrins to mediate binding to odontoblasts, monocytes, and endothelial cells (de Vega *et al*, 2007; Ikeuchi *et al*, 2018; Sarangi *et al*, 2018; Tsunezumi *et al*, 2018). However, the role of fibulin 7 in the skin remains largely unknown.

*Fbln7* knockout (KO) mice did not show apparent changes in skin histology or epidermal thickness (Fig. S3A–C), but a modest trend toward increased proliferation in the 2- to 3-month-old *Fbln7* KO scale IFE was observed, as indicated by BrdU labeling (Fig. S3D, E, F). To test whether fibulin 7 modulates epidermal stem cell heterogeneity and behavior during aging, we applied lineage tracing tools and labeled fast-cycling stem cells with Slc1a3^CreER^ in the *Fbln7* KO background. Fast-cycling stem cell clones were localized in the scale region at 1 week or 3 months post-labeling, and some clones persisted for up to 1 year in *Fbln7* wild-type (WT) mice (Fig. 3A [upper panel], B, C). In contrast, after 1 year, the number and size of fast-cycling stem cell clones in the scale region of *Fbln7* KO mice were significantly decreased compared with *Fbln7* WT mice (Fig. 3A [lower panel], B, C). Negative impacts on fast-cycling stem cells were not observed at earlier time points, indicating the age-dependent function of *Fbln7* in the longterm maintenance of the fast-cycling stem cell population (Fig. 3A–C). Next, we attempted to compare the slow-cycling stem cells labeled with Dlx1^CreER^ in WT and *Fbln7* KO mice; however, due to the proximity of the *Dlx1* promoter to the *Fbln7* gene locus on chromosome 2, we could not obtain *Fbln7* KO mice harboring Dlx1^CreER^. Comparisons between *Fbln7* WT and *Fbln7* heterozygous (Het) mice from 1-week, 3-month, and 1-year chases revealed that Dlx1 clones were enriched in the interscale region and that their quantity and distribution showed no differences between WT and Het mice (Fig. S4A, B). Similarly, a smaller number of Slc1a3 clones in the interscale and their respective clonal areas were not affected by the loss of *Fbln7* (Fig. S4C, D), suggesting that *Fbln7* may not affect slow-cycling stem cells in the interscale.

**Figure 3.**
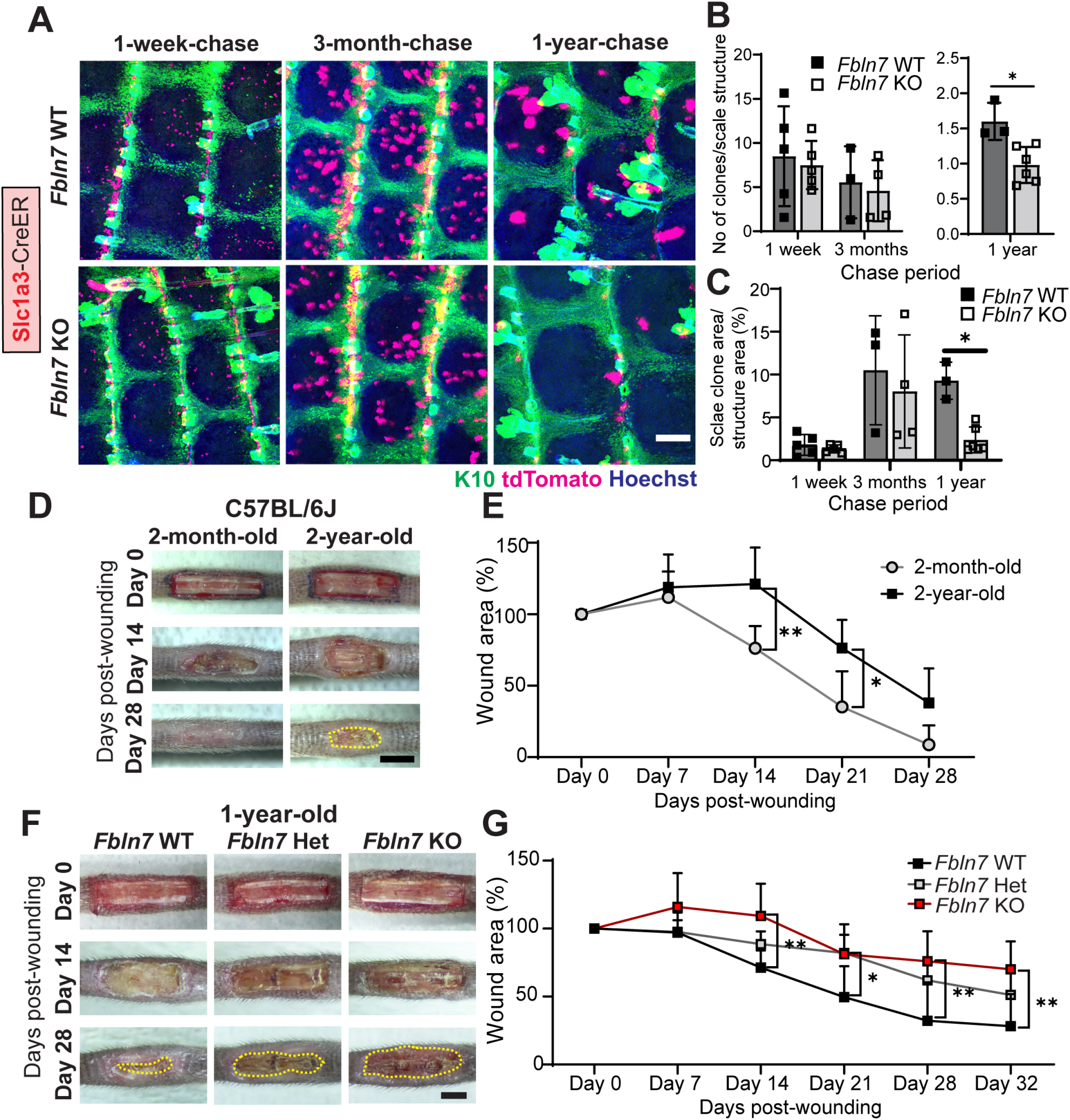
*Fbln7* knockout leads to aggravated fast-cycling stem cell loss and impaired injury repair. **A.** Slc1a3^CreER^ lineage tracing in the *Fbln7* WT and KO backgrounds. Low-dose tamoxifen was administered once at 2 months of age and samples were collected after 1-week, 3-month, and 1-year chases. Wholemount staining of tail epidermis with tdTomato, K10, and Hoechst. Scale bar 200 μm. **B.** The number of Slc1a3^CreER+^ clones per scale structure. **C.** Quantitation of the area of Slc1a3^CreER+^ clones per total area of scale structure. *Fbln7* WT mice for 1-week (*N*=5), 3-month (*N*=3), and 1-year chases (*N*=3). *Fbln7* KO mice for 1-week (*N*=5), 3-month (*N*=4), and 1-year chases (*N*=6). (B, C) Mann–Whitney test. **D.** Representative pictures from tail wounds of 2-month vs. 2-year-old C57BL/6J wild type mice. Scale bar 4 mm. **E.** Measurements of wound area over time in 2-month vs. 2-year-old C57BL/6J mice, *N*=6 and *N*=5, respectively. **F.** Representative pictures from tail wounds of 1-year-old *Fbln7* WT, heterozygotes (het), and KO mice. Scale bar, 4 mm. **G.** Measurements of wound area over time in *Fbln7* WT (*N*=5), het (*N*=6), and KO (*N*=7) mice. All graphs in Figure 3 show mean ± S.D. *; *P*<0.05. **; *P*<0.01. Statistical tests for (E, G) were performed with two-way ANOVA, Tukey’s multiple comparisons test. All graphs show mean ± S.D.

Slow- and fast-cycling stem cells work together in response to skin damage (Sada *et al*., 2016), and wound healing is delayed in aging skin (Keyes et al., 2016; Liu et al., 2019). Our experiments also confirmed that wound healing was significantly delayed in 2-year-old mice compared to 2-month-old mice by measuring the speed of wound healing in the tail skin of C57BL/6J wild-type mice (Fig. 3D, E). We hypothesized that disturbed epidermal stem cell heterogeneity in chronological aging or the absence of *Fbln7* might reduce skin regenerative capacity upon injury. To test this possibility, a wound-healing assay was performed in the tail skin of 2- to 3-month-old and 1-year-old mice in the presence or absence of *Fbln7* (Fig. 3F–G, S4E, F). While no apparent difference was observed in the wound healing ability of *Fbln7* WT or KO mice at 2–3 months of age (Fig. S4E, F), 1-year-old *Fbln7* KO mice showed impaired wound closure compared with *Fbln7* WT mice (Fig. 3F, G). These results suggest that the expression of *Fbln7* in aging skin may minimize the loss of epidermal stem cell heterogeneity, supporting resilience against damage to skin tissue.

### Fibulin 7 modulates stem cell inflammatory stress response and fate specification

To further understand how fibulin 7 supports long-term stem cell maintenance, RNA-seq was performed using basal epidermal stem cells isolated from the dorsal skin of *Fbln7* WT and KO mice at 2 months or 1 year of age. The dorsal skin was utilized as it is the largest skin area containing the slow- and fast-cycling stem cell populations in the IFE (Sada *et al*., 2016). PCA analysis showed that the transcriptomic difference between *Fbln7* WT and KO mice occurred at 1 year but not at 2 months of age, supporting our previous findings from lineage tracing and wound healing assays (Fig. 4A, Table S3). Intriguingly, in the 1-year-old *Fbln7* KO mice, inflammatory response genes were upregulated, including those involved in antigen presentation, the MAPK cascade, and cytokine production, while chemotaxis genes were downregulated (Fig. 4B-D, Table S3). Intriguingly, in the 1-year-old *Fbln7* KO mice, inflammatory response genes were upregulated, including antigen presentation, the MAPK cascade, and cytokine production while chemotaxis was downregulated (Fig. 4B-D, Table S3). Lineage fate changes have been reported as part of the inflammatory response (Ge *et al*., 2017), and an analysis of these gene lists (Ge *et al*., 2020) further underscored their alteration in 1-year-old *Fbln7* KO mice (Fig. 4E). The changes resemble some features of the 2-year-old nLRCs in the tail (Fig. 2G), such as enhanced hair lineage and IFE differentiation markers, albeit with some mouse-to-mouse variations (Fig. 4E).

**Figure 4.**
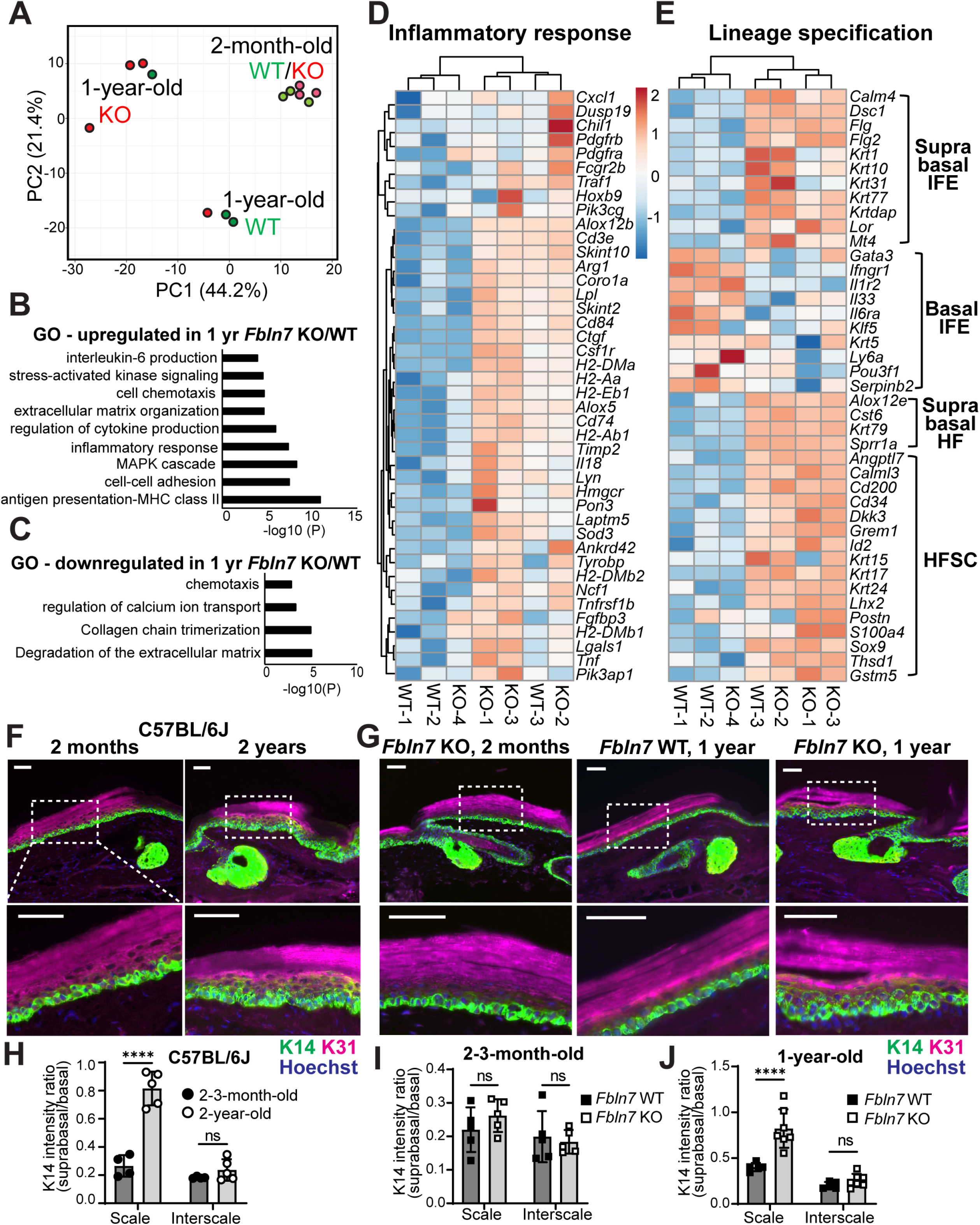
Loss of *Fbln7* is associated with an increased inflammatory response in aging epidermal stem cells and lineage misregulation. **A.** PCA map constructed from 3-fold differentially expressed genes in epidermal stem cells from *Fbln7* WT vs. KO dorsal skin in 2-month- and 1-year-old mice. Each dot represents one mouse (*N*=3 per group, except 1-year-old *Fbln7* KO *N*=4). **B, C.** Gene ontology analysis from ≥2-fold upregulated (B) or downregulated (C) genes in 1-year-old *Fbln7* KO mice compared with WT. **D.** Heatmap illustrating changes in inflammatory response genes in 1-year-old *Fbln7* WT vs. KO mice, constructed from ≥2-fold differentially expressed genes obtained from gene ontology analysis in (B). **E.** Heatmap describing changes in basal and suprabasal signature genes of epidermal stem cells and HFSCs (Ge *et al*., 2020) in 1-year-old *Fbln7* WT vs. KO mice. Heatmaps scale show log2 RPKM values. **F, G.** K31 and K14 immunostaining in tail sections of 2-month- and 2-year-old C57BL/6J wildtype mice (F) and 1-year-old *Fbln7* WT vs. KO compared with 2-month-old *Fbln7* KO mice (G). Dotted box areas are enlarged in the lower panels. Scale bars, 50 μm. Hoechst, nuclear staining. **H.** Quantification of K14 intensity in the suprabasal area normalized to basal area per cell. Each dot represents an average value from one mouse. *N*=4 for 2-3-month-old and *N*=5 for 2-year-old C57BL/6J mice. **I, J.** K14 intensity quantification as (H) performed in 2-3-month-old and 1-year-old *Fbln7* WT vs. KO mice. *N*=5 for both WT and KO of 2-3-month-old mice. *N*=5 and *N*=7 for 1-year-old WT vs. KO, respectively. All statistical tests in (H-J) were performed with two-way ANOVA, Tukey’s multiple comparisons test. All graphs show mean ± S.D. ****; *P*<0.0001, ns; not significant.

Increased inflammation due to DNA damage has been linked to stem cell fate misspecification, which is manifested in the suprabasal expression of basal marker K14 (Seldin & Macara, 2020). We tested whether the slow- and fast-cycling stem cells of the tail epidermis of 1-year-old *Fbln7* KO or 2-year-old C57BL/6J WT mice also undergo lineage misspecification by examining the distribution of K14 expression. K14 was restricted to the basal layer in 2-month-old C57BL/6J WT mice, as opposed to the basal-to-suprabasal expansion of K14 observed primarily in the scale region of the 2-year-old counterparts (Fig. 4F). Likewise, there was an increased tendency for basal-to-suprabasal expansion of K14 in the scale region of the 1-year-old *Fbln7* KO mice but not in the 2-month-old *Fbln7* KO mice (Fig. 4G). Quantifications confirmed that the increase in suprabasal K14 was significant in the scale but not in the interscale (Fig. 4H-J). Although suprabasal K14 was not observed in aged dorsal skin (Keyes *et al*., 2016), it has been reported that aging promotes an inflammatory environment in skin (Doles *et al*, 2012; Hu *et al*, 2017); fast-cycling stem cells in the tail scale may thus be more susceptible to this change. Therefore, our results indicate that *Fbln7* loss increased the expression of inflammatory response genes in epidermal stem cells, which may, in turn, influence the specification of stem cell fate.

### Fibulin 7 maintains the extracellular environment of epidermal stem cells by physically interacting with matrix proteins

In the 1-year-old *Fbln7* KO mice, in addition to inflammatory response genes, ECM-related genes were also affected, with changes impacting basement membrane (BM) components (collagen IV, nidogen 1, laminins, fibulin 1) and ECM remodeling genes (Fig. 4B, C, 5A). This suggests that the loss of fibulin 7 may modify ECM composition in the stem cell microenvironment. To further characterize the biochemical functions of fibulin 7, we screened for secreted fibulin 7-binding proteins using conditioned media prepared from heparin-treated cells overexpressing full-length (FL) fibulin 7 or cells overexpressing fibulin 7 but lacking the N-terminal heparin-binding coiled-coil domain (dCC) (Tsunezumi *et al*., 2018). The putative fibulin 7-binding proteins were co-eluted with fibulin 7 from a metal ion affinity column and identified via mass spectrometry (Fig. S5A). The following classes of candidate proteins were selected: those with functions in the structural integrity of the BM, growth factors, or proliferation-modulating proteins; matricellular proteins associated with wound healing; and extracellular proteases involved in ECM remodeling (Fig. S5B). We confirmed via solid-phase binding assay the dose-dependent interactions of fibulin 7 with BM components such as collagen IV and, to a lesser extent, the fibulin 1C and 1D isoforms (Fig. 5B, C).

**Figure 5.**
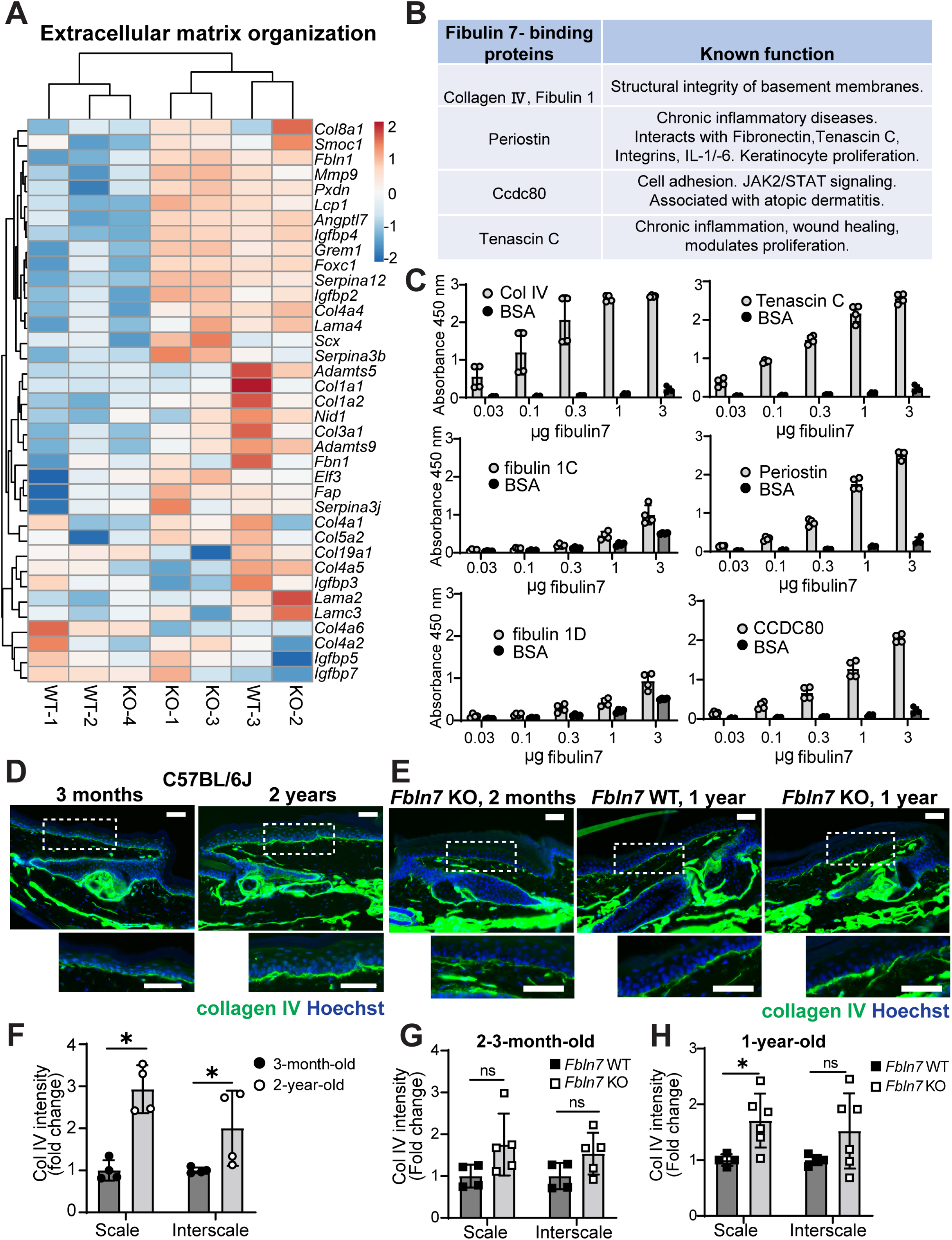
Transcriptome of *Fbln7* KO mice and fibulin 7-binding assays indicate fibulin 7 mechanism of action through ECM regulation. **A.** Heatmap of ≥2-fold upregulated or downregulated genes related to ECM from gene ontology analysis in figure 4B–C. Scale reflects log2 RPKM values. **B.** Shortlisted fibulin 7-binding protein candidates (from Figure S6A, B) and their reported functions. **C.** Solid-phase binding assays using recombinant fibulin 7 as liquid phase and purified or recombinant ECM proteins as solid phase. X-axis shows increasing doses of fibulin 7 (μg/well). Bovine serum albumin (BSA) was used as the control liquid phase and added in the same amounts as fibulin 7. Graph shows means ± S.D. (4 technical repeats in 2 independent experiments). **D, E.** Collagen IV immunostaining in tail sections of 3-months-old vs 2-year-old C57BL/6J mice (D) and 1-year-old *Fbln7* WT vs. KO compared with 2-month-old *Fbln7* KO mice (E). Dotted box regions are enlarged. Scale bars, 50 μm. Hoechst, nuclear staining. **F, G, H.** Quantification of Collagen IV basement membrane intensity per cell in 2-year-old mice normalized to 3-month-old mice in C57BL/6J mice (F) or in *Fbln7* KO mice normalized to WT mice (G, H). Graphs show mean ± S.D. Mann-Whitney test. *; *P*<0.05, ns; not significant.

The structural proteins of the BM are essential in maintaining epidermal stem cells in physiological and pathological conditions (Abreu-Velez & Howard, 2012; Chermnykh *et al*, 2018; Gatseva *et al*, 2019; Watanabe *et al*., 2017). Collagen IV staining illustrated an uneven thickening pattern along the BM of 2-year-old versus 3-month-old C57BL/6J mice (Fig. 5D). A similar pattern was observed in the epidermis of 1-year-old *Fbln7* KO mice compared to 1-year-old WT or 2-month-old *Fbln7* KO mice (Fig. 5E). This aligns with a previous report describing the uneven distribution of collagen IV in 2-year-old epidermal BM (Changarathil *et al*, 2019) and upregulation of collagen IV in thickened BM, which led to niche stiffness in aged skin (Koester *et al*., 2021). Quantifications of BM collagen IV intensity demonstrated a significant increase in the scales of 2-year-old C57BL/6J and 1-year-old *Fbln7* KO epidermis, but not in 2–3-month-old *Fbln7* KO mice (Fig. 5F-H). In the interscale, BM collagen IV staining showed a similar increase during chronological aging (Fig. 5F), though this increase was not statistically significant in the *Fbln7* KO mice (Fig. 5G, H). Moreover, fibulin 7 exhibited direct binding to matricellular proteins that are upregulated upon wound healing or inflammation, such as tenascin C, periostin, and Ccdc80 (Fig. 5B, C) (Hirota *et al*, 2012; Midwood *et al*, 2016; Nikoloudaki *et al*, 2020; Tremblay *et al*, 2009), but not to insulin growth factor binding protein-2 (IGFBP-2, Fig. S5C). These results suggest that fibulin 7 supports epidermal stem cells through physical interactions with ECM proteins, thereby maintaining the extracellular environment of the skin.

### Fibulin 7 regulates the undifferentiated state and proliferative response to growth factors in primary keratinocytes

In vivo phenotype analysis has so far suggested that fibulin 7 may function as a protective factor for epidermal stem cells during physiological aging, maintaining their long-term stem cell potential. To investigate the function of fibulin 7 in primary mouse keratinocytes, we overexpressed the FL or dCC mutant (Fig. 6A–C). The CC domain mediates binding to heparin and is important for the pericellular tethering of fibulin 7 (Tsunezumi *et al*., 2018). Overexpression of both FL fibulin 7 and the dCC mutant suppressed differentiation markers (*Krt1, Krt10, Lor, Dsc1*), whereas stem cell markers (*Krt14, Klf5*) and cobblestone-like morphology of keratinocytes were unchanged, suggesting that fibulin 7 maintains primary keratinocytes in an undifferentiated state (Fig. 6A, D) independently of its CC domain.

**Figure 6.**
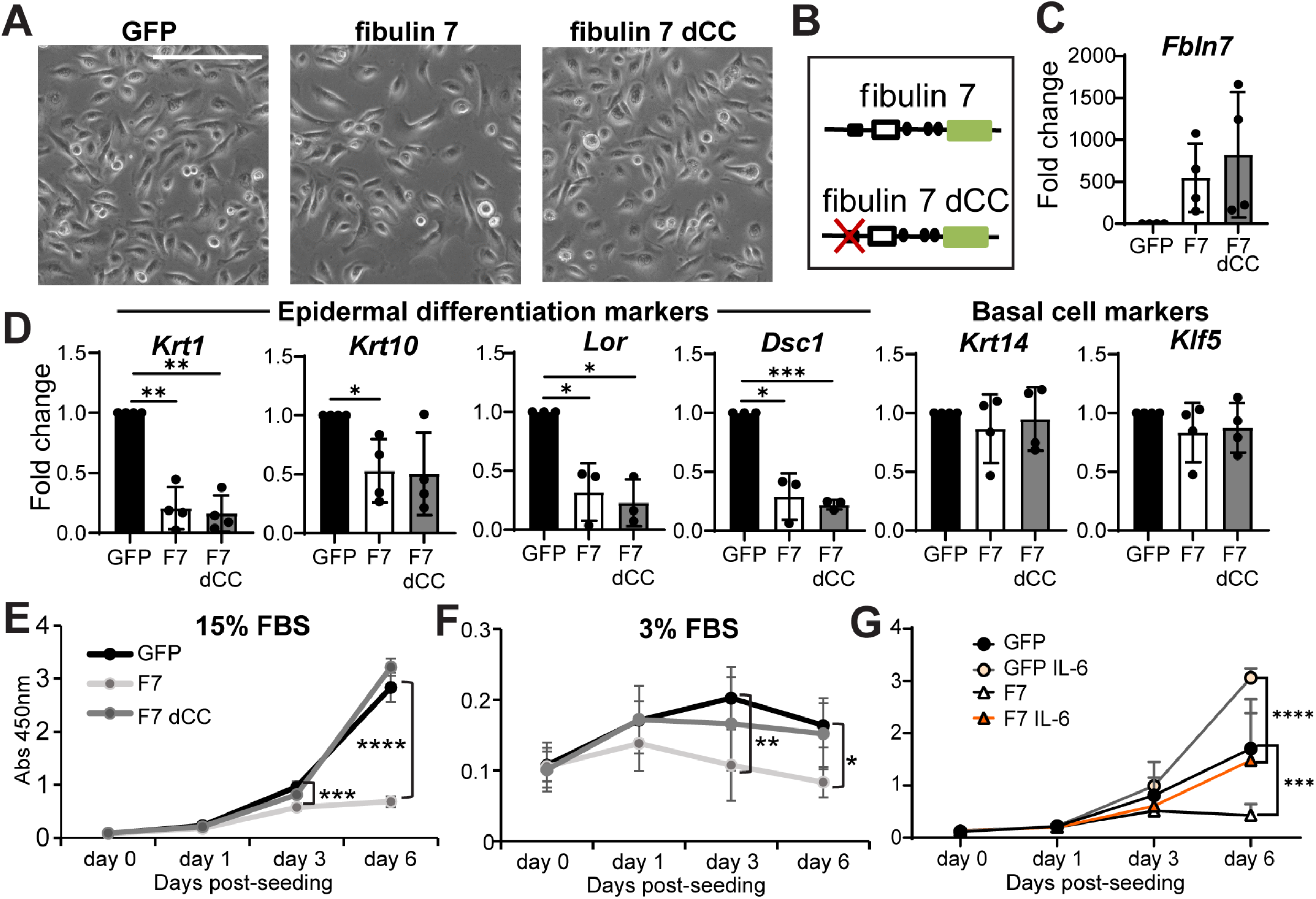
Fibulin 7 overexpression maintains the undifferentiated, slower proliferation state and confers resistance to IL-6-induced proliferation in primary keratinocytes. **A, B.** Brightfield images of mouse primary keratinocytes overexpressing GFP, full-length fibulin 7 (F7), or coiled-coil deletion mutant (F7 dCC) as shown in (B). Scale bar, 200 μm. **C, D.** Quantitative RT-PCR shows fold changes in gene expression of F7 or F7 dCC compared with GFP control. *N* = at least 3 biological repeats. Welch’s t-test. ***; *P*<0.001. **; *P*<0.01. *; *P*<0.05. Mean ± S.D. **E, F.** Cell proliferation assays comparing growth rates of GFP overexpression control with F7 and F7 dCC over 6 days in 15% FBS growth medium (E) or 3% reduced growth medium (F). **G.** Cell proliferation assay comparing GFP or F7 overexpressing keratinocytes treated with 5 ng/ml IL-6 over 6 days. Graphs in (E-G) describe mean absorbance values ± S.D. from at least 3 biological repeats and statistically tested with two-way ANOVA (Tukey’s multiple comparisons test). ****; *P*<0.0001.

We further investigated the role of fibulin 7 in keratinocytes exposed to a high or low amount of serum growth factors, mimicking the aging skin microenvironment. The proliferation assays in high (15%) or low (3%) serum conditions demonstrated that fibulin 7 maintains a lower cell division frequency of keratinocytes and that this effect required the CC domain (Fig. 6E, F). FL fibulin 7 overexpression also inhibited expression of the *Igfbp2* gene, which promotes stem cell proliferation (Huynh *et al*, 2011; Ichijo *et al*, 2017) (Fig. S5D). GO analysis from 1-year-old *Fbln7* KO RNA-seq data indicated an upregulation of genes associated with interleukin-6 (IL-6) production (Fig. 4B). IL-6 induces hyperproliferation of keratinocytes in vitro and is associated with skin aging, wounding, and psoriasis (Barrientos *et al*, 2008; Doles *et al*., 2012; Grossman *et al*, 1989; Hu *et al*., 2017; Taniguchi *et al*, 2014). We found that responses to growth cues by this cytokine were blunted in fibulin 7-overexpressing keratinocytes (Fig. 6G). Thus, fibulin 7 gain-of-function indicates that fibulin 7 may be beneficial for suppressing differentiation and maintaining slower proliferation status in epidermal stem cells under high or low growth-inducing conditions.

## Discussion

Young skin is supported by heterogeneous stem cell populations having the high capacity for injury recovery and resilience to stress. The current study highlights the importance of epidermal stem cell heterogeneity and its dysregulation during physiological aging (Fig. 7). We demonstrated that the Slc1a3+ fast-cycling epidermal stem cell clones were decreased and the cellular and molecular properties of epidermal stem cells impaired in aged skin. The deletion of *Fbln7*, an ECM gene, accelerated the age-dependent loss of fast-cycling stem cell clones, accompanied by defective skin regeneration after tissue damage. Fibulin 7 maintains epidermal stem cells in part by regulating inflammatory stress responses and the fate balance of stem cells through direct binding to ECM proteins. Our results suggest that regulation of the extracellular environment by fibulin 7 maintains heterogeneous epidermal stem cell populations over the long term, which may be an essential mechanism in controlling skin aging.

**Figure 7.**
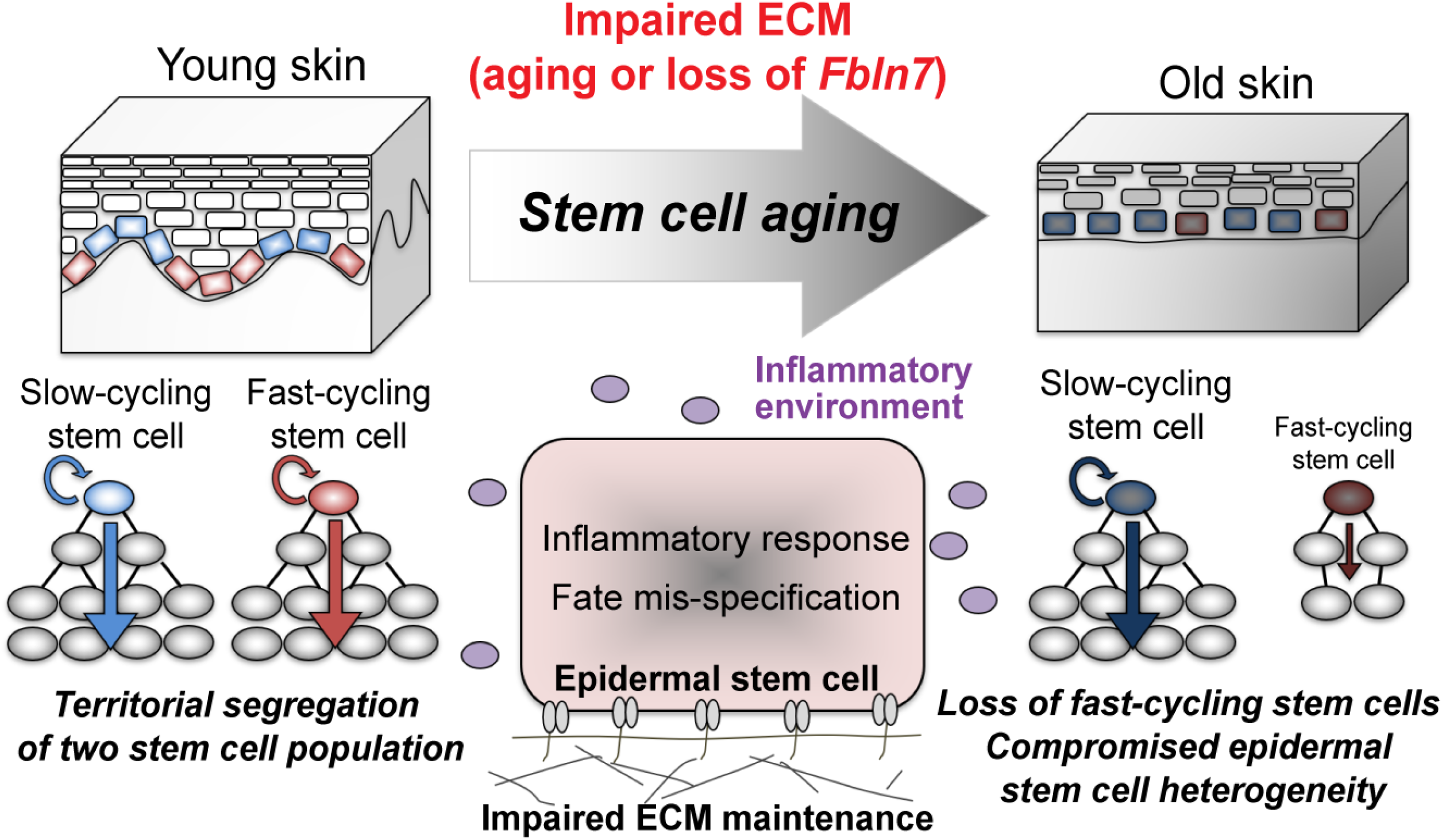
Proposed model. Fibulin 7, a matricellular protein, is necessary for long term maintenance of epidermal stem cells heterogeneity and skin regenerative capacity by supporting ECM organization and stem cell response to aging-related inflammation.

Loss of stem cell polyclonality and positive selection for clones with driver mutations have been observed in aging and are closely linked to carcinogenesis (Martincorena *et al*, 2015; McKerrell & Vassiliou, 2015). Since fast-cycling stem cells may accumulate more replication-induced mutations over time, the loss of fast-cycling clones could be advantageous for suppressing tumorigenesis in aged skin. The Slc1a3 population may have a low competitive advantage, as it is particularly prone to loss in the scale region compared to the Col17a1 population (Liu *et al*., 2019). However, while aged slow-cycling clones in the interscale were likely to have long-term stem cell potential, the impaired expression of DNA damage repair genes (Fig. 2C) may correlate to the incidence of basal cell carcinoma in the aging population (Nagarajan *et al*, 2019; Sanchez-Danes *et al*, 2016; Youssef *et al*, 2010). Nonetheless, DNA-damaged stem cells could be eliminated from tissues via differentiation (Kato *et al*, 2021), which may account for the relatively low incidence of cancer compared to the rate of mutations occurring (Martincorena *et al*., 2015). Further study is needed to determine whether there are differences in DNA damage repair mechanisms or the behavior of DNA-damaged cells between slow- and fast-cycling stem cell populations.

ECM composition is modified tremendously during skin aging, an alteration that is partially caused by ECM gene expression changes (Table S1) (Ge *et al*., 2020; Koester *et al*., 2021; Liu *et al*., 2019; Marsh *et al*, 2018; McCabe *et al*, 2020). *Fbln7* expression was increased in fast-cycling stem cells persisting in 2-year-old mice, and *Fbln7* KO exacerbated the loss of this stem cell population and impaired wound healing ability (Fig. 3). RNA-seq data confirmed the age-dependent effects of fibulin 7 loss at 1 year of age, which intriguingly indicated an increased inflammatory response, together with augmented suprabasal differentiation markers and HFSC lineage fate abnormalities, as observed in the 2-year-old mice (Fig. 4). This suggests that fibulin 7 may be upregulated in aged skin to protect epidermal stem cells from the adverse consequences of chronic inflammation, such as inhibition of self-renewal and skewed stem cell lineages (Doles *et al*., 2012; Pietras *et al*, 2016). A recent study suggests that IFE and hair follicle BMs are thickened with age, and protein components of the BM, such as laminins and collagen IV, are upregulated, contributing to stiffening of the stem cell niche and decreased stem cell function (Koester *et al*., 2021). Future studies should evaluate whether the binding of fibulin 7 to BM proteins (Fig. 5) supports BM structural integrity and remodeling under physiological and pathological conditions of the skin. The binding of fibulin 7 to β1 integrin (de Vega *et al*., 2007; Ikeuchi *et al*., 2018; Sarangi *et al*., 2018) could also be relevant in maintaining epidermal stem cells. β1 integrin plays an immuno-protective role by inhibiting inflammation-induced ECM degradation in the BM (Kurbet *et al*., 2016), and its loss results in IFE proliferation and differentiation aberrations, BM abnormalities, and defective injury repair (Lopez-Rovira *et al*, 2005; Margadant *et al*, 2010; Watt, 2002). It is worth noting that the reported IFE changes due to β1 integrin loss (Lopez-Rovira *et al*., 2005) bear some similarities to our description of aging tail skin, such as K10-marked interscale expansion (Fig. 1E, F) and increased collagen IV deposition in the BM (Fig. 5).

Overexpression of fibulin 7 slowed proliferation and enhanced the undifferentiated state of primary keratinocytes (Fig. 6), in contrast to the differentiation-biased state caused by loss of fibulin 7 (Fig. 4). Our in vivo data, however, showed no significant changes in epidermal stem cell proliferation in *Fbln7* KO mice (Fig. S3). Compensatory actions by other factors like IGFBP2, which is suppressed in fibulin 7-overexpressing cells and increased in 1-year-old *Fbln7* KO epidermal stem cells, may explain these differences (Fig. 5A, Fig. S6D). Of note, the slower cycling rate induced by fibulin 7 overexpression was only partially enhanced by IL-6, which is a positive regulator of keratinocyte proliferation (Taniguchi *et al*., 2014). This indicates that fibulin 7 may dampen keratinocyte response to excess inflammatory cytokine signaling. Fibulin 7 could represent a potential molecule for protecting epidermal stem cells against the detrimental effects of aging-associated inflammation, lineage misspecifications, and a premature differentiation-like state. Pending future research, fibulin 7 could be a valuable candidate for the intervention of chronic wounds observed more frequently in older patients.

## Limitations of the study

Since basal cells in the scale proliferate less than those in the interscale (Changarathil *et al*., 2019; Giangreco *et al*, 2008), separation by cell division frequency in the H2B-GFP system became more ambiguous in aged mice and may not exactly represent the same stem cell populations in the young mice. Furthermore, the loss of Slc1a3^CreER^+ fast-cycling stem cell clones during aging made it difficult to isolate the same stem cell population labeled at younger ages. Therefore, although our interpretation of RNA-seq data largely reflects molecular changes that have occurred in slow- and fast-cycling stem cells, in-depth analysis by single-cell analysis or isolation using specific cell surface markers will advance the understanding of epidermal stem cell aging.

In our RNA-seq analysis, we used tail and back skin which had distinct gene expression profiles and responses to inflammation (Quigley *et al*, 2016) due to different originating skin locations. Although the effects of aging may differ between dorsal skin (Ge *et al*., 2020; Keyes *et al*., 2016) and tail skin, it is worth noting that *Fbln7* KO back skin showed molecular changes (Fig. 4E) comparable to those of aging tail skin (Fig. 2G). In the future, RNA-seq using tail skin from *Fbln7* KO will allow for direct comparison with physiological aging.

Finally, since we use a whole-body knockout of *Fbln7*, we cannot definitively determine if *Fbln7* functions primarily in the dermis or immune cells rather than epidermal stem cells. The in vivo detection of fibulin 7 protein in mouse tissues has been challenging. Although experiments with keratinocytes support the intrinsic effects of *Fbln7*, phenotype analysis using conditional knockout mice is needed to investigate the cell type-specific functions of *Fbln7*.

## Methods

### Mice

All animal experiments were performed according to the Institutional Animal Experiment Committee guidelines at the University of Tsukuba and Kumamoto University. The generation of *Fbln7* KO mice (129SvEv;C57BL/6J) was previously reported (Tsunezumi *et al*., 2018). K5-tTA (C57BL/6J) (Diamond *et al*, 2000)/pTRE-H2B-GFP (CD1;C57BL/6J) (Tumbar *et al*., 2004) double transgenic mice were used for the isolation of LRCs and non-LRCs from young and old mice (Sada *et al*., 2016). Wild-type C57BL/6J mice were purchased from Charles River Laboratories or Japan SLC, Inc. For the lineage-tracing experiment, Dlx1^CreER^ (C57BL/6J) (The Jackson Laboratory, no. 014551) or Slc1a3^CreER^ (C57BL/6J) (The Jackson Laboratory, no. 012586) mice were crossed with Rosa-tdTomato reporter mice (C57BL/6J) (The Jackson Laboratory, no. 007905). CreER/Rosa-tdTomato was introduced in the *Fbln7* knockout background. Mice of both sexes were used for experiments.

### Tamoxifen, BrdU, and doxycycline administration

Tamoxifen (Sigma) was injected intraperitoneally at a single dose of 25 μg or 50 μg/g body weight (BW) of Slc1a3^CreER^ and Dlx1^CreER^ mice, respectively, at 2 months of age for lineage tracing experiments. BrdU (5-Bromo-2’-deoxyuridine; Sigma-Aldrich, B5002) was administered in drinking water (0.8 mg/ml) two days before sacrifice to label proliferative cells. For H2B-GFP pulse-chase, 2-month-old or 22-month-old mice were fed with doxycycline chow (1 g/kg, Oriental Kobo Inc.) for two weeks before sacrifice.

### FACS isolation

The subcutaneous and fat tissues were removed from the skin and incubated overnight at 4 °C in 0.25% trypsin/EDTA and the next day at 37 °C for 30 minutes. Single-cell suspensions were prepared by gentle scraping of the epidermis and subsequent filtering with 70 μm and 40 μm filters. Cells were stained with the following antibodies for 30 minutes on ice: CD34-biotin (1:50, eBioscience, 13-0341), Streptavidin-APC (1:100, BD Biosciences, 554067), α6-integrin-BUV395 (1:100, BD Biosciences, custom order), and Sca1-BV421 (1:100, BD Biosciences, 562729). Dead cells were excluded by propidium iodide (P4864, Sigma) staining. Cells from mouse tail skin (aging RNA-seq) and dorsal/ventral skin (fibulin 7 RNA-seq) were isolated using FACS Aria flow cytometer (BD Biosciences). Data were analyzed using FlowJo software (BD Biosciences).

### RNA-sequencing

FACS-isolated cells from K5-tTA/pTRE-H2B-GFP mice and *Fbln7* WT and KO mice were directly sorted into Trizol (Ambion, 10296028) and submitted to Tsukuba i-Laboratory LLP, the University of Tsukuba, for further analysis. RNA integrity was analyzed with Agilent 2100 bioanalyzer. The RNA-seq library was prepared with SMARTer^®^ Stranded Total RNA-Seq Kit v2 – Pico Input Mammalian (Takara), and sequencing was performed in the Next Generation Sequencing platform using NextSeq500 (Illumina). The data were analyzed with CLC genomics 11 software (Qiagen). After normalization, genes that showed ≥2-fold changes were selected, and hierarchical clustering and gene ontology analysis were performed using the online tools VENNY 2.1 (Oliveros, 2007–2015; https://bioinfogp.cnb.csic.es/tools/venny/index.html), Morpheus (https://software.broadinstitute.org/morpheus), Clustvis, Panther (www.geneontology.org)(Ashburner *et al*, 2000; Gene Ontology, 2021; Mi *et al*, 2019a), and Metascape (Metsalu & Vilo, 2015; Mi *et al*, 2019b; Zhou *et al*, 2019).

### Wholemount and section immunostaining of mouse tail epidermis

Tail skin wholemount and frozen section staining procedures were performed as previously described (Sada *et al*., 2016). The primary antibodies and dilutions used included the following: rat anti-BrdU (1:300, Abcam, ab6326), rabbit anti-K14 (1:1000, BioLegend, 905304), mouse anti-K10 (1:100, BioLegend, 904301, or 1:100, Abcam, ab9026), guinea pig anti-K31 (1:100, PROGEN Biotechnik, GP-hHa1), rabbit anti-collagen IV (1:200, Millipore, AB756P) and chicken anti-GFP (1:500, AB13970). Mouse on Mouse (M.O.M) kit reagents (Vector Laboratories) were added for blocking when a primary mouse antibody was used. Secondary antibodies (Alexa 488, 546, 647, Invitrogen) were diluted at 1:200. All samples were counterstained with Hoechst (Sigma, B2261) before mounting. Before anti-BrdU antibody incubation, samples were treated with 2N HCl at 37 °C for 1 hour. The stained wholemount epidermis was observed under a confocal microscope (Zeiss LSM 700) and images were captured and analyzed using ZEN 2010 software. All wholemount pictures are shown as projected Z-stack images, viewed from the basal side. A Zeiss Axio Imager Z2 fluorescence microscope was utilized for stained sections and images were acquired with Zen 2.3 Pro software with final adjustments in Adobe Photoshop.

### Hematoxylin and eosin (H&E) staining

Tail skin was directly embedded in optimal cutting temperature (OCT) compound (Tissue-Tek, Sakura). Ten μm sections were fixed in 4% paraformaldehyde (PFA) at room temperature for 10 minutes. Sections were stained with hematoxylin (Wako, 131-09665) for 20 minutes and eosin Y (Wako, 058-00062) for 15 seconds before dehydration and mounting in Entellan solution (Merck Millipore, HX73846161). Images were captured using a Zeiss Axio Imager Z2 microscope and Zen 2.3 Pro software.

### Quantification and statistical analysis

All quantifications were independently performed on at least 3 mice. Data are shown as mean ± standard deviation (S.D). In H&E-stained tail sections, an epidermal unit was defined as the IFE region comprising 1 scale and 1 interscale structure, often between 2 hair follicles. The epidermal thickness was quantified from 6 epidermal units per mouse (Changarathil *et al*., 2019). The scale and interscale IFE thicknesses were measured at the center of an epidermal unit and the IFE region adjacent to the hair follicle, respectively. K10/K31 positive areas, the number of tdTomato positive cell clones, clonal areas, and BrdU+ cells were manually scored from 4–8 images consisting of 16–32 interscale or scale IFE structures per mouse in projected Z-stack images using ImageJ software (NIH). Using the same software, K14 intensity ratio between suprabasal/basal cells and Collagen IV basement membrane intensity were measured from 5 cells/structure/picture and 4 pictures were used to obtain an average score per mouse. Statistical tests are described in the respective figure legends.

### Wound healing assay

Full-thickness tail wounding was performed as described previously at 10 mm length × 3 mm width on the dorsal side of the tail (Liu *et al*., 2019). Ketoprofen (2 μg/g BW, Sigma) and amoxicillin (100 μg/g BW, Sigma) were intraperitoneally injected prior to tail cutting. After wound pressurization to stop bleeding, film spray dressing (Cavilon, 3M) was applied to the wound surface, and mice were monitored until recovery. To follow up on wound size over time, images were taken with a Zeiss Stemi305 microscope with an Axiocam208 camera every 7 or 8 days until day 32 post-wounding, and the wound area was measured by ImageJ.

### Lentivirus induction of *Fbln7* overexpression in primary mouse keratinocytes

Isolation of 2-day-old C57BL/6J mouse primary keratinocytes was performed as previously reported (Lichti *et al*, 2008). Keratinocytes were cultured in low-calcium (0.05 mM) E-medium containing 15% chelex-treated Fetal Bovine Serum (FBS) and used for experiments from passage 7–15. Cell images were acquired using an Evos FL cell imaging system (Thermo Fischer Scientific).

Mouse FL or dCC mutant *Fbln7* with V5-His tag (Tsunezumi *et al*., 2018) was subcloned into a CSII-CMV-MCS-IRES2-Bsd vector (RIKEN, Tsukuba, Japan). The CSII-CMV-MCS-IRES2-Bsd vector (*Fbln7* FL, dCC or control *eGFP*) was transfected into 293T cells using polyethylenimine reagent (PEI, Polysciences Inc.) together with lentivirus packaging vectors (pRSV-Rev, pMD2.G, and pMDLg/pRRE) (Addgene). Lentivirus-containing medium was concentrated using Lenti-X-concentrator (Takara Bio). Six-well plates were seeded with 100,000 primary keratinocytes per well and transduced with 150 uL each of lenti-eGFP, lenti-*Fbln7*, or lenti-*Fbln7* dCC and 4 μg/mL polybrene (Sigma). The medium was changed the next day, and cells were cultured for 2-3 days before selection with blasticidin at a concentration of 1 μg/mL.

### Cell proliferation assay

Primary keratinocytes were seeded at 2000 cells/well in 96-well plates with technical duplicates or triplicates. Experiments were repeated three times unless indicated otherwise in the figure legend. Cell proliferation was measured using Cell Counting Kit-8 (CCK-8, Dojindo) as previously described (Oinam *et al*, 2020). Recombinant mouse interleukin-6 (R&D Systems, 406-ML) was added to the culture medium at the indicated concentration.

### Quantitative RT-PCR

RNA was isolated using RNeasy mini kit (Qiagen). cDNA was synthesized using iScript cDNA synthesis kit. RT-PCR was performed using iTaq Universal SYBR green supermix (Bio-Rad). Mouse primer sequences are as follows: *Klf5* forward (F) 5’-GGCTCTCCCCGAGTTCACTA-3’ and reverse (R) 5’-ATTACTGCCGTCTGGTTTGTC-3’; *Krt14* F 5’-AAGGTCATGGATGTGCACGAT-3’ and R 5’-CAGCATGTAGCAGCTTTAGTTCTTG-3’; *Krt1* F 5’-AACCCGGACCCAAAACTTAG-3’ and R 5’-CCGTGACTGGTCACTCTTCA-3’; *Krt10* F 5’-GGAGGGTAAAATCAAGGAGTGGTA-3’ and R 5’-TCAATCTGCAGCAGCACGTT-3’; *Lor* F 5’-TCACTCATCTTCCCTGGTGCTT-3’ and R 5’-GTCTTTCCACAACCCACAGGA-3’; *Dsc1* F 5’-GGGAGCACCTTCTCTAAGCA-3’ and R 5’-CACTCTTCCAGATTCACTTTGCC-3’; *Igfbp2* F 5’-CGCTACGCTGCTATCCCAAC-3’ and R 5’-TCGTCATCACTGTCTGCAACC-3’; *Fbln7* F 5’-GAGGAGGCTTCCAGTGTGTC-3’ and R 5’-AATGGAAGGAGATGGTCTTGG-3’; *Gapdh* F 5’-TGCCCAGAACATCATCCCT-3’ and R 5’-GGTCCTCAGTGTAGCCCAAG-3’.

### Metal ion affinity chromatography and mass spectrometry

Mouse fibulin 7 (*Fbln7*) coding sequence (Tsunezumi *et al*., 2018) or human *FBLN7* lacking the coiled-coil domain (ΔCC) (Primers: F 5’-GAACTGTCCAGATGCCCTTCCAGTT-3’ and R 5’-GCATCTGGACAGTTCTGGGAAGCCCG-3’) was cloned by PCR and ligated into the pEF6/V5-His TOPO plasmid vector (Invitrogen). FBLN7-V5-His vector was transfected into CHO cells (FBLN7-CHO) and ΔCC-V5-His vector was transfected into HEK293T cells (ΔCC-293T) to express fibulin 7 as a COOH-terminally V5-His-tagged recombinant protein. Both cell types were maintained in a mixture of Dulbecco’s Modified Eagle’s medium and Ham’s F-12 medium (DMEM/F12) (Invitrogen) supplemented with 10% FBS.

FBLN7-CHO cells were grown to confluence and the resulting serum-free conditioned medium (CM) was harvested every 2 days. The collected CM was concentrated by protein precipitation with 80%-saturated ammonium sulfate (Sigma) and dissolved in phosphate-buffered saline without calcium and magnesium [PBS (-)] supplemented with 0.1% Triton X-100 and a proteinase inhibitor mixture [0.2 mM 4-(2-aminoethyl) benzenesulfonyl fluoride (AEBSF), 0.16 μM aprotinin, 0.025 mM bestatin, 7.5 μM E-64, 0.01 mM leupeptin, and 5 μM pepstatin] (Calbiochem). The precipitants were dialyzed against a 400 times volume of PBS (-) plus Triton X-100 and then applied to a Nickel (Ni)-Sepharose Histidine-affinity column (His GraviTrap; GE Healthcare). Bound proteins were eluted with PBS (-) supplemented with 500 mM NaCl, 500 mM imidazole, and 0.01% Triton X-100. To exclude non-specific bound proteins, the fractions containing FBLN7 were applied to a heparin-Sepharose-6B column (GE Healthcare). FBLN7 bound to the column and mainly eluted at 0.6 M NaCl. This fraction was further purified by Ni-Sepharose column chromatography and the final elute fraction containing FBLN7 and its putative high-affinity binding proteins was subjected to SDS-PAGE. Separated proteins were detected by staining with Coomassie Brilliant Blue R250.

293T and ΔCC-293T cells were grown to confluence and the resulting serum-free conditioned medium (CM) was harvested every 2 days. The collected CM was concentrated by protein precipitation with 80%-saturated ammonium sulfate (Sigma) and dissolved in 50 mM Tris-HCl (pH 7.4) containing 150 mm NaCl supplemented with 5 mM CaCl_2_, 0.1% Triton X-100, and a proteinase inhibitor mixture (as previously described). The precipitants were dialyzed against a 400 times volume of diluent buffer. The dialyzed 293T CM was passed through a Cobalt-affinity column (ThermoFisher Scientific) once to eliminate non-specific bound proteins, and the flow-through fraction was collected and mixed with ΔCC, and then applied to the column chromatography. ΔCC and its affinity-binding proteins bound to the column, and 1200 mM NaCl was used to elute the proteins that were associated with ΔCC. The eluted samples were subjected to SDS-PAGE and separated proteins were visualized by Coomassie staining as previously described. The excised proteins from both preparations were submitted to the Proteomics Core at the University of Texas Southwestern Medical Center for analysis.

### Solid-phase binding assay

Ninety-six-well flat-bottom MaxiSorp Nunc plates (Invitrogen) were coated with 1 μg/well of recombinant mouse periostin, Ccdc80 (R&D Systems), mouse fibulin 1C, fibulin 1D, human collagen IV (Sasaki *et al*, 1995), human tenascin C (Matsui *et al*, 2018), or BSA controls (Sigma) in bicarbonate buffer (15 mM Na_2_CO_3_, 35 mM NaHCO_3_, pH 9.2) at 4 °C for 16 hours. All incubations thereafter were performed at room temperature. After 3 washes with Ca^2+^-Mg^2+^ free Dulbecco Phosphate Buffered Saline (DPBS, Gibco), wells were blocked with 5% non-fat dry milk in Tris Buffered Saline (TBS) for 1 hour. As a liquid phase, 0.03 to 3 μg/well of the soluble mouse or human HA-tagged recombinant fibulin 7 (R&D Systems) in 2% dry milk/TBS and 2 mM CaCl_2_ were added to the wells and incubated for 3 hours. Wells were then washed 5 times with wash buffer (TBS, 0.025% Tween-20, 2 mM CaCl_2_). Anti-HA antibody (3724S, Cell Signaling Technology) was diluted 1/1500 in 2% dry milk TBS buffer and added to the wells for 1.5 hours. After 5 washes, secondary anti-rabbit antibody (170-6515, Bio-Rad) was applied at 1/1000 dilution for 1 hour, followed by another 5 washes. Colorimetric reaction was performed using Substrate Reagent Pack (DY999, R&D Systems) according to the manufacturer’s protocol and absorbance was read at 450 nm using an xMark microplate reader (Bio-Rad).

## Data availability

RNA-sequencing data is deposited in the Gene expression omnibus (GEO) under accession number, GSE185086 and GSE185087. All raw mass spectrometry data files have been deposited to the Mass spectrometry Interactive Virtual Environment (MassIVE; Center for Computational Mass Spectrometry at the University of California, San Diego) and can be accessed using the MassIVE ID MSV000088142.

## Acknowledgments

We thank the Animal Resource Center at the University of Tsukuba and the Center for Animal Resources and Development at Kumamoto University for their excellent mouse care, M. Higashi and T. Keida for technical help and M. Goodarzi and A. Lemoff and the Proteomics Core Laboratory at the University of Texas Southwestern Medical Center for assistance with proteomics data analysis. We would like to thank Dr. M. Kato, Dr. H. Suzuki, and Dr. Y. Watanabe (University of Tsukuba) for their help in lentivirus production. We also thank Dr. T. Suda (Kumamoto University) for critical reading of this manuscript and Dr. M. Muratani (University of Tsukuba) for his advice on RNA-seq analysis. This work was supported by AMED-PRIME, AMED (JP21gm6110016) (to A.S.), AMED under Grant Number 21bm0704067 (to A.S.), Grant-in-Aid for Scientific Research (B) (20H03266) (to A.S.), Grant-in-Aid for Research Activity Start-up (20K22659) (to E.R.), Grant-in-Aid for Scientific Research on Innovative Areas “Stem Cell Aging and Disease” (17H05631) (to A.S.), Grant-in-Aid for Research Activity Start-up (16H06660) (to A.S.), and Grant-in-Aid for Early-Career Scientists (18K14709) (to A.S.). This work was supported by the following research grants: The American Heart Association (12EIA8190000), The Mizutani Foundation for Glycoscience, The Uehara Memorial Foundation, and The Tokyo Biochemical Research Foundation to H.Y., and The Naito Foundation, The Nakajima Foundation, The Astellas Foundation for Research on Metabolic Disorders, The Uehara Memorial Foundation, The Inamori Foundation, The Takeda Science Foundation, The Koyanagi Foundation, The Nakatomi Foundation, Leave a Nest Co., Ltd. for IKEDARIKA award, and Basic Research Support Program Type A, University of Tsukuba (all to A.S.), and the Cooperative Research Project Program of Life Science Center for Survival Dynamics, Tsukuba Advanced Research Alliance (TARA Center) (to A.S. and J.T.). This work was also supported by JST SPRING, Grant Number JPMJSP2124 (to Y.X.N).

## Author contributions

A.S., H.Y., and E.R. conceptualized the study, designed the experiments, and wrote the manuscript. G.C. and E.R. performed and analyzed the lineage tracing and RNA sequence experiments. L.O., E.R., and R.I. performed and analyzed the *Fbln7* knockout mouse experiments. E.R., J.T., L.O., T.S., and K.I. performed and analyzed the fibulin 7 binding assays. E.R. and Y.X.N. performed and analyzed the keratinocyte culture experiments. A.S. and H.Y. supervised the project. A.S., H.Y., and E.R. acquired funding. All authors provided input on the final manuscript.

